# Proteolytic Performance is Dependent on Binding Efficiency, Processivity and Turnover: Single Protease Insights

**DOI:** 10.1101/2024.06.10.598230

**Authors:** Emily Winther Sørensen, Freya Björk Reinhold, Andreas Faber, Steen Bender, Jacob Kaestel-Hansen, Jeannette de Sparra Lundin, Errika Voutyritsa, Per Hedegaard, Sune M. Christensen, Nikos S. Hatzakis

## Abstract

Proteases are essential enzymes for a plethora of biological processes and biotechnological applications, e.g., within the dairy, pharmaceutical, and detergent industries. Decoding the molecular level mechanisms that drive protease performance is key to designing improved biosolutions. However, direct dynamic assessment of the fundamental partial reactions of substrate binding and activity has proven a challenge with conventional ensemble approaches. We developed a single-molecule (SM) assay for the direct and parallel recording of the stochastic binding interaction of Savinase, a serine-type protease broadly employed in biotechnology, with casein synchronously with monitoring proteolytic degradation of the substrate. SM recordings enabled us to determine how the overall activity of Savinase and two mutants relies on binding efficiency, enzymatic turnover and activity per binding event. Analysis of residence times revealed three characteristic binding states. Mutations were found to dominantly alter the likelihood of sampling the long lived state, with lifetimes longer than 30 seconds, indicating this state contributes to overall activity and supporting a level of processivity for Savinase. This observation challenges conventional expectations, as the protease has no characterized substrate binding site, or binding domain, aside from the active site. These insights, inaccessible through conventional assays, offer new perspectives for engineering proteases with improved hydrolytic performance.

## INTRODUCTION

Proteases hydrolyse the peptide bonds in proteins^1,2^, making them important catalysts in living organisms and invaluable in biotechnological industries^3–6^. Savinase is an alkaline serine endo-protease belonging to the subtilisin superfamily^7–9^. It plays a paramount role in biotechnology due to its high stability and broad substrate specificity and is widely applied for protein-based stain-removal in detergency^10^ as well as in, e.g., protein extraction^11^ and biocatalysis^12^. Thus, leads on how to further engineer Savinase, and protease reactions more broadly, can open an avenue to greener biosolutions.

While we have an overall understanding of protease reactions^13,14^, the precise steps and interactions involved in substrate binding and degradation are still subject to ongoing research^7,15–17^. The reactions of biotechnological interest are almost exclusively interfacial, and adsorption of the protease to the substrate surface is a fundamental requirement for activity. The surface environment in turn impacts the substrate encounter rate (i.e binding efficiency) to the substrate and thus hydrolytic performance (proteolytic degradation of substrate)^10^. This is, e.g., evident in a study of a chymotrypsin serine protease interacting with a protein multilayer that demonstrated the importance of electrostatics in driving protease adsorption but also revealed a non-trivial dependence of the activity (i.e. performance) on the density of protease bound to the insoluble substrate^18,19^. Interestingly, in that system, a fraction of stably recruited proteases exhibited mobility at the surface, which could explain some of the observed variations in specific activity of the studied charged variants. The impact of the surface environment is also evident for Savinase that shows a different activity profile in heterogeneous versus homogeneous reaction conditions^10^. Curiously, Savinase has been shown to exhibit conformational dynamics in the region around the active site which may have functional significance on substrate binding^20,21^. The evidence emphasizes the importance of further studies to elucidate protease function in relevant interfacial contexts^10,18,19^ but also the need for new tools to cope with spatiotemporal complexities of molecular mechanisms. While providing priceless insights to the field, previous studies averaged the function of a large ensemble of proteases, masking how proteolytic performance is dependent on SM binding efficiency, enzymatic turnover and activity per binding event.

Here, we report a single-molecule (SM) imaging assay^22–28^ to directly observe Savinase binding and degradation of proteinaceous casein micelles^29^ at the fundamental limit of individual proteases. Casein, a protein found in milk^30^, is an important protease substrate and its proteolytic degradation is essential in the dairy industry for the production of cheese^31^. As such, casein is regularly used in enzyme performance testing^32^. The developed assay enabled us to determine how the overall performance of three Savinase variants depends on the binding efficiency and dynamics, activity per binding event and turnover of bound proteases. To the best of our knowledge, this is the first example of an assay capable of simultaneously monitoring dynamic binding behavior and substrate degradation of proteases on a SM level.

We found that the differing performance of two of the three variants examined was readily explained by one variant exhibiting a higher number of binding events, i.e. higher binding efficiency, to the micellar substrate. However, a variant with intermediary performance displayed the least efficient substrate association. Despite its inferior binding efficiency, this variant exhibited the highest activity per binding event and the largest relative turnover number, suggesting processivity of Savinase - per se an unexpected characteristic for an enzyme without a known binding site or domain distinct from the active site^8,33^. The residence times of all transient binding events revealed the existence of three characteristic dwell-times, inferred as distinct binding states of Savinase at the casein micelles. Notably, we find residence times in the second scale with a long-lived binding state exceeding 30 seconds and we show that sampling of the binding states is sensitive to mutations in the protease. The demonstration that Savinase is capable of lingering at the substrate surface at timescales, far exceeding expectations for single hydrolytic turnovers per substrate surface association, corroborates the notion of processivity, i.e., multiple turnovers per binding event, and as such it further supports that the protease reaction mechanism has commonalities with classical interfacial enzyme reactions^34–37^.

## RESULTS AND DISCUSSION

### Direct Observation of Substrate Degradation Synchronously with Single Protease Binding Behavior in Real-Time

We developed a Total Internal Reflection Fluorescence (TIRF) microscopy-based assay enabling direct observation of individual Savinase molecules interacting with surface immobilized casein substrate (Figure 1A). Specific and singular labeling of Savinase was achieved by employing single cysteine mutants labeled with SeTau647-maleimide. Substrate degradation was monitored in real-time by employing Alexa Fluor 488 labeled casein that forms large protein micelles (Figure 1B) in the pH range of 7-12^29^, where Savinase has the highest activity^8^. The formation of casein micelles creates a heterogeneous size distribution of substrate^29^ and ensures that most casein micelles have multiple fluorescent labels (Figure S1). Enzymatic release of labeled casein-derived peptides by Savinase leads to their diffusion into solution and the concomitant decrease in fluorescence provides a measure for proteolytic performance (proteolytic degradation and removal of casein from the surface) (Figure 1B). Dual color imaging and our quantitative image analysis^38–41^ allowed the parallel recording of the stochastic and reversible binding of Savinase to the casein covered surface and its degradation (Figure 1B/C).

**Figure 1.**
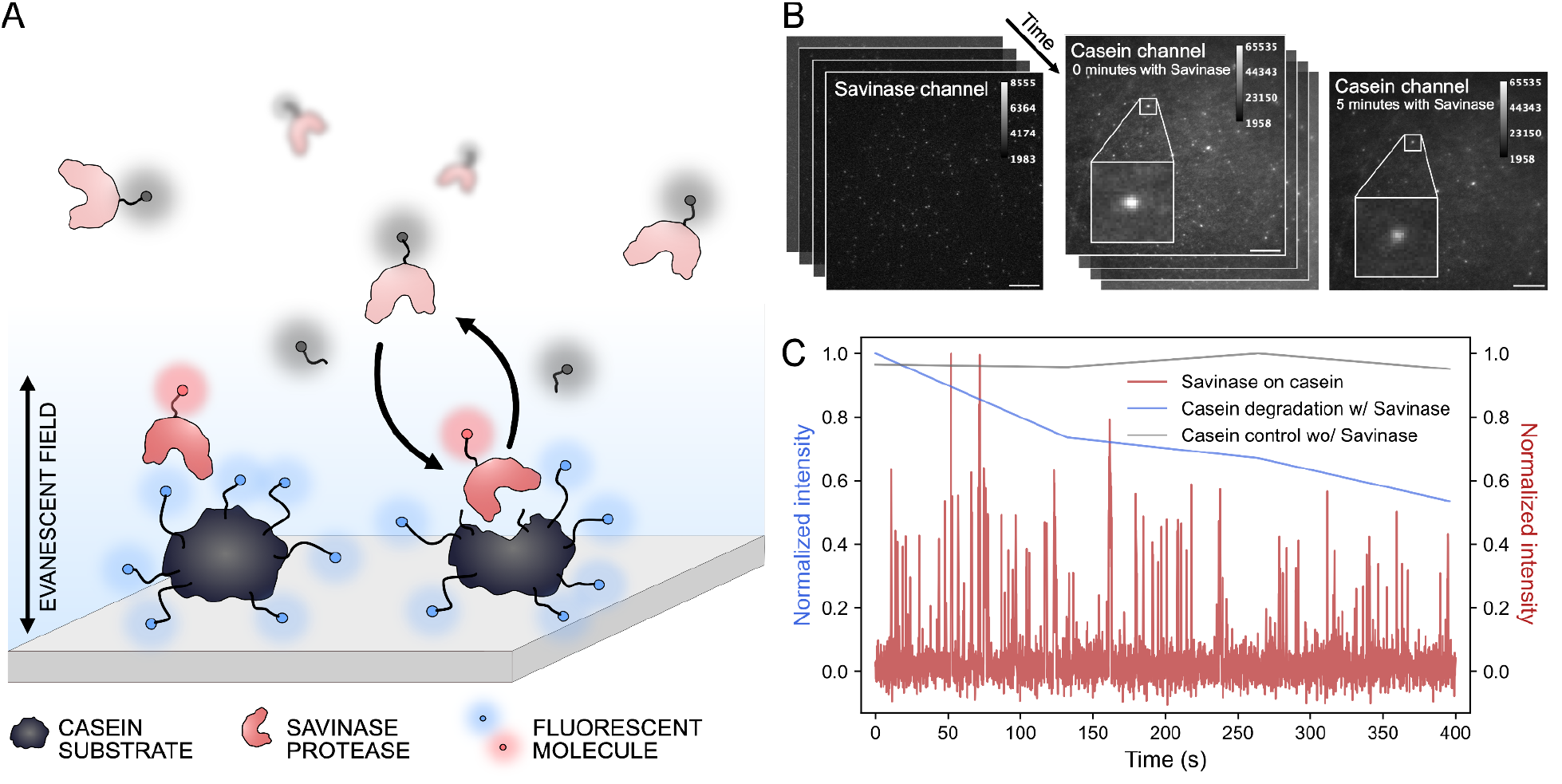
Experimental setup for direct observation of casein substrate degradation from transient binding of individual Savinase proteases. A) Representation of experimental setup for real-time recordings of the reversible binding of individual SeTau647-labeled Savinase proteases on Alexa488-labeled casein substrate synchronously with substrate degradation. The evanescent field allows for recording of only proteases in proximity to the casein covered surface, not in solution. Upon proteolytic cleavage of casein the cleaved fluorescently labeled peptide chains diffuse in solution resulting in signal loss in the casein channel. B) From left: Representative experimentally acquired time series of 0.05 nM Savinase on casein covered surface. Center: Time series of the casein channel showing the direct observation of individual casein micelles. Right: representative frame recorded 6 minutes after the addition of Savinase, displaying visible signal loss in casein channel, highlighted by zoom of same casein micelle in center figure. Scale bar is 10 μm for all images and the calibration bar indicates intensity scale of gray value^42^. C) Representative normalized intensity traces, of sum of intensity in region of interest (ROI) around casein micelle, displaying the synchronous recording of the reversible binding of Savinase proteases (red trace) on a single casein micelle and the intensity loss of that casein micelle (blue trace) due to proteolytic cleavage and diffusion of fluorescent peptide in solution. Note the different temporal resolution in imaging to avoid casein signal photobleaching. Grey trace shows the control casein trace without Savinase showing minimal bleaching within the experimental time frame.

At the utilized concentration of casein, a dense layer of substrate micelles is deposited on the glass surface (Figure S2). Savinase is found to show little to no difference in binding behavior to large casein micelles as compared to small (Figure S3). Thus, we accumulate data from the entire field of view in the analysis of binding behavior and performance. To minimize photobleaching and maximize output we recorded 1 frame in the casein channel, per 1000 frames in the Savinase channel. For each surface, a total of 4004 frames were recorded at 10 Hz and the experimental time was approximately 7 minutes. As such, the assay allows for parallelized recording of thousands of individual Savinase binding events at each casein surface.

### Correlation of SM Binding Efficiency and Performance of Savinase Variants Indicates Processivity

We employed the assay to study three protease variants: wild-type Savinase (S1), Savinase 2 (S2) and Savinase 3 (S3), all bearing a single exposed cysteine at position 135 for specific labeling (see Methods and Materials). The mutations in S2 and S3 reside in the periphery of the cleft down to the active site and result in a more negative net charge of S2 and S3 as compared to S1. Specifically, S2 carries an arginine to serine mutation, as compared to S1, while S3 features an insertion of aspartic acid (see Figure S4). The relative performance of the variants measured by the SM assay is validated by its agreement with corresponding ensemble performance measurements as seen in Figure 2A (see Methods and Materials and Figure S5), where S1 was found to have the highest performance and S2 the lowest. Comparative analysis of the three variants establishes a basis for deconvoluting Savinase performance in the casein system.

**Figure 2.**
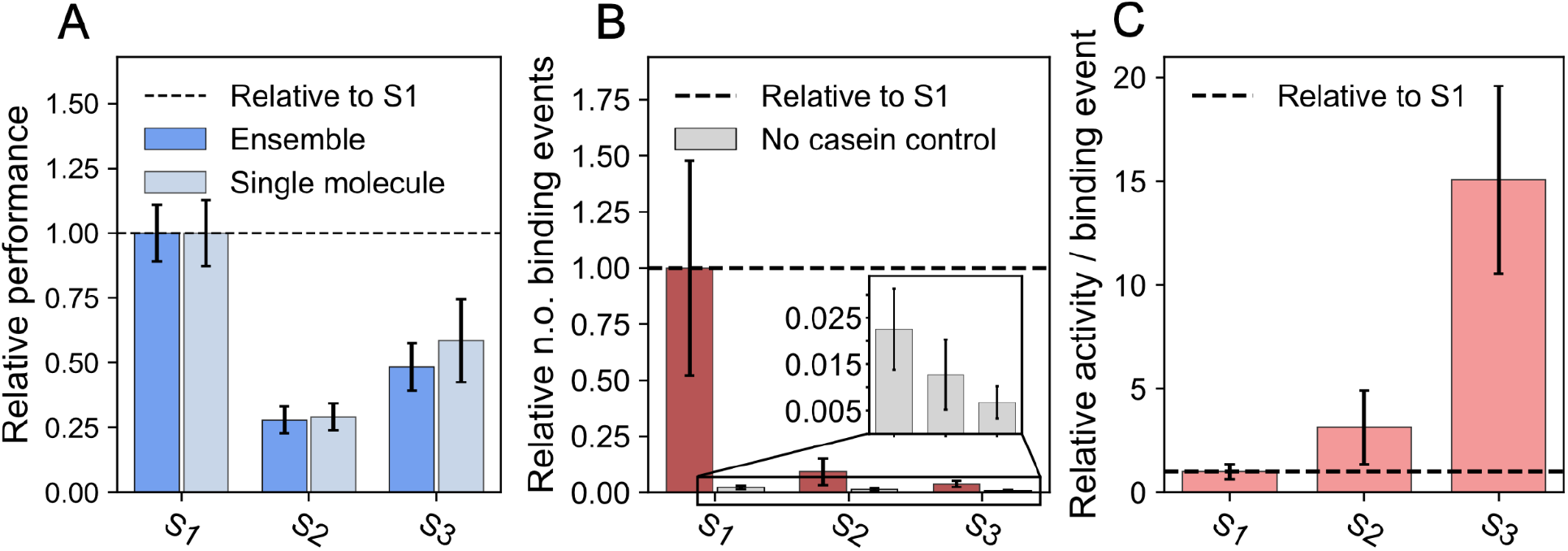
Performance, binding efficiency and processivity of three Savinase variants. A) Relative performance of three Savinase variants, S1, S2 and S3, on a SM and ensemble basis (see Figure S4). S1 variant is 3.4 and 1.7 times more active than S2 and S3 respectively. B) Relative quantification of number of Savinase binding events, the binding efficiency, for each of the three variants on casein within the experimental time frame (see Table S2 for absolute numbers). S1 variant displays an 11 and 26 times higher binding efficiency than S2 and S3, respectively. C) Relative activity per binding event, i.e. processivity, of Savinase variants, namely the performance (A) divided by the recorded number of binding events (B). S2 and S3 display 5 and 15 times more activity per binding event, i.e. processivity, respectively, than S1. Error bars correspond to the standard deviation of three technical replicates.

The SM readout allows the extraction of the number of reversible Savinase binding events on the casein substrate over the course of experimental time, i.e., the binding efficiency (Figure 2B). S1 displayed the highest binding efficiency and S3 the lowest. This is consistent with the expected net charge of S2 and S3 being more negative than S1, which likely results in an increased repulsion to the negatively charged casein at pH 8^43^, supporting the influence of electrostatic interactions on protease substrate^44,45^. However, inspection of Figure 2A and B reveals that the overall performance ranking of the Savinase variants cannot be explained solely by variations in binding efficiency.

While the high performance of S1 is consistent with S1 showing the highest binding efficiency among the three variants, S3 has intermittent performance as compared to S1 and S2, but displays the lowest number of binding events. This reveals a difference in activity on the substrate, where S2 and S3 exhibit higher activity per binding event than S1 (Figure 2C). These data demonstrate that the overall system-level performance is driven both by the number of proteases bound as well as the activity of individual bound proteases. A corollary is that multiple cleavage events can occur within a single binding event, i.e., processivity^33^. In principle, it could also be that not all binding events are productive but, as we shall see below, it is unlikely to account for our observations given the lifetime of these interactions. It is intriguing that Savinase may catalyze multiple peptide bond cleavages within a single binding event, despite lacking known substrate binding sites beyond its active site^8^. However, it is qualitatively in alignment with the previous report on chymotrypsin protease^18,19^ suggesting this could be a recurring feature of interfacial protease reactions.

### Residence Times of Savinase on Substrate Reveals Three Binding States

Analysis of residence time for thousands of detected reversible SM binding events on the casein covered surface provided further insight on the three Savinase variants (Figure 3A/B and S6). The residence time distributions for the Savinase variants were broad and showed an exponentially decaying behavior^46^ (Figure 3C and S3). Unbinned maximum likelihood estimate (MLE)^47^ showed optimal fitting by a mixture model of the weighted sum of three exponential functions (equation 1), indicating the existence of three distinct binding states of Savinase. In this context, a binding state refers to a temporary condition where Savinase engages with the casein substrate with a characteristic dwell-time. The probability distribution is given in equation 1, where P(*t*_i_) is the probability density of Savinase binding for a duration of t_i_ and W_1_, W_2_ and W_3_ are the weights of the exponential fit segment with a lifetime of *τ*_1_, *τ*_2_ and *τ*_3_, respectively.

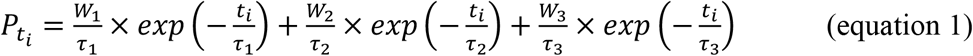

**Figure 3.**
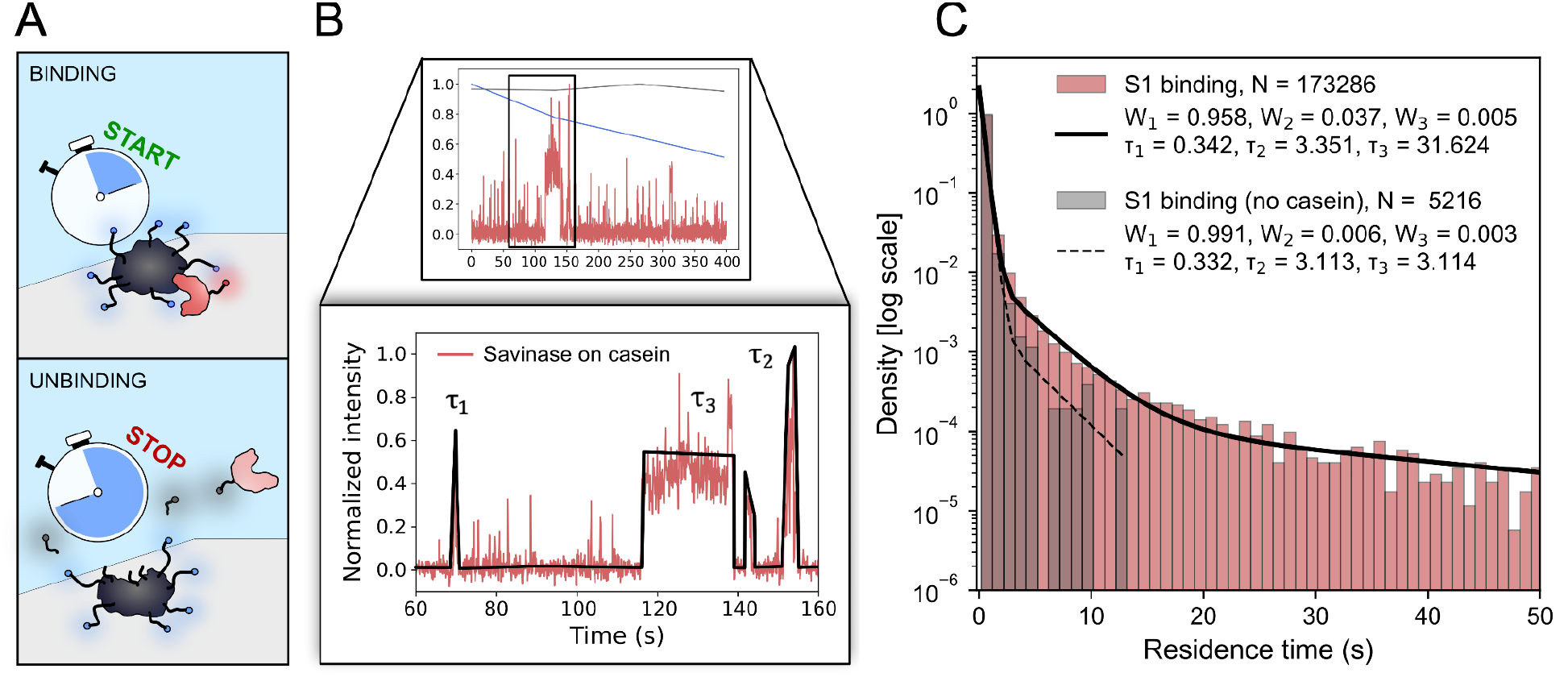
Extraction of residence times of all bound Savinase molecules reveals three underlying binding behaviors. A) Representation of how the binding duration of each Savinase molecule that binds to the substrate was extracted from the recorded videos. B) Representative trace displaying the reversible and stochastic binding of individual S1 molecules on single casein micelle. Zoom in displays intensity spikes corresponding to the signal of the reversible binding of S1 molecules of different durations, shown qualitatively by the black trace. C) Probability distribution of residence times of all bound S1 molecules of four replicates, with (red) and without (gray) casein substrate. The distributions were optimally fitted with a triple exponential distribution (equation 1) using unbinned MLE (see Methods and Materials), indicating the existence of three binding states for S1, which persisted for S2 and S3. In the absence of casein Savinase residence times can be fit with the first two of the three states sampled in the presence of casein (see Figure S8 for S2 and S3 distributions and fits, with and without casein).

We can consider the weight of each exponential segment as the probability of sampling a given binding state, or the relative number of Savinase molecules represented in that binding state, where the lifetimes indicate the duration of binding. Fitting of residence times for S1, S2 and S3, show the existence of 1) a highly occupied short-lived state with a lifetime *τ*_1_ ∼0.5 seconds accounting for 80-95% of all bound Savinase molecules, W_1_; 2) an intermediate-lived state with a lifetime *τ*_2_ ∼3-5 seconds, accounting for 4-15% of the bound molecules, W_2_; and 3) a long-lived scarcely occupied state with a lifetime, *τ*_3_ ∼30-60 seconds, only accounting for 0.5-5% of all bound molecules, W_3_ (Table 1). Although the long-lived state was less frequently sampled, it is substantial in the time domain and it is highly sensitive to mutations, suggesting functional significance. For S1, the long-lived state accounts for ca. 20% of the total time proteases were observed in contact with the casein covered surface whereas for S2 and S3 it accounts for more than 60% (Figure S7 and Table S2).

**Table 1.** Occupancy and duration of the three binding states for three Savinase variants. Table summarizing the number of binding events, and the occupancy and duration of the three extracted binding states for the three variants, S1, S2 and S3, in the presence of casein (see Figure S8 for no casein control). All errors represent the standard deviation of three technical replicates.

|  | S1 w/ casein | S2 w/ casein | S3 w/ casein |
| --- | --- | --- | --- |
| **N** | $57762 \pm 19527$ | $5368 \pm 2878$ | $2240 \pm 277$ |
| **Sampling of binding states (% / N)** |  |  |  |
| **W1** | $95.84 \pm 0.47 /$<br>$55361 \pm 18717$ | $81.16 \pm 0.66 /$<br>$4357 \pm 2336$ | $87.12 \pm 1.47 /$<br>$1952 \pm 243$ |
| **W2** | $3.69 \pm 0.46 /$<br>$2131 \pm 767$ | $14.79 \pm 1.20 /$<br>$793 \pm 431$ | $10.25 \pm 1.34 /$<br>$230 \pm 41$ |
| **W3** | $0.47 \pm 0.03 /$<br>$270 \pm 93$ | $4.05 \pm 0.65 /$<br>$218 \pm 122$ | $2.62 \pm 0.43 /$<br>$59 \pm 12$ |
| **Duration of binding states (seconds)** |  |  |  |
| **$\tau_1$** | $0.34 \pm 0.01$ | $0.44 \pm 0.04$ | $0.45 \pm 0.02$ |
| **$\tau_2$** | $3.35 \pm 0.10$ | $4.15 \pm 0.36$ | $5.17 \pm 0.62$ |
| **$\tau_3$** | $31.62 \pm 4.23$ | $54.72 \pm 12.04$ | $61.18 \pm 11.18$ |

Control experiments conducted without casein on the surface substantiate these results. Firstly, the total number of binding events decreased significantly, respectively, 44, 7 and 6 fold for S1, S2 and S3 (Table S2), showing the majority of binding events observed on the substrate-covered surfaces pertains to the protease-casein interaction. Secondly, the long-lived binding state, *τ*_3_, was not observed in the data without casein (Figure 3C). Thus, the third and long-lived binding state is only sampled in the presence of substrate, strongly supporting it being an attribute of protease activity. Analysis of subsets of the recorded residence times to account for potential under-sampling further confirmed the finding of a long-lived (≈30-60 s) binding state of the protease specific to casein surface (Figure S9).

Comparison of the three binding states between the Savinase variants shows that although S1 is binding to the surface more frequently (Figure 2B), it is rarely sampling the longer-lived states, W_2_ and W_3_. S1 also exhibits the shortest longer-lived states, *τ*_2_ and *τ*_3_ (Table 1). Thus, the fact that S1 exhibits the lowest processivity (Figure 2C), implies that in addition to being proteolytically active, longer-lived states may facilitate greater degradation by proteases. The existence of long-lived active states further supports the idea of Savinase being processive. This observation challenges conventional understanding and suggests that Savinase may have structural features that allow it to sustain its interaction with the substrate for multiple cleavage events, despite the absence of well-defined binding domains. While these features might have an electrostatic nature^18,19^, the observation of diminished binding efficiency despite the higher net negative charge of S2 and S3 relative to S1 suggests a more complex underlying phenomenology. Interestingly, our data indicate that this ability is highly sensitive to mutations. Some enzymes, such as lipases, demonstrate processivity by traversing the substrate surface^34–36^. In contrast, Savinase seems to stay anchored locally on the casein layer within the timeframe of a binding event. One explanation could be that its processivity might arise from digging into the substrate rather than relocating laterally.

### Parameterizing performance

The direct recordings of reversible binding events allowed us to decipher the performance generating molecular mechanisms in terms of 1) the binding efficiency, 2) processivity (activity per binding event) and 3) turnover number (Figure 4 and 5). The turnover number (Figure 4) is given by the relative number of cleavage events per time, calculated as the ratio of the overall performance (Figure 2A) and the sum of residence times for all observed binding events.

**Figure 4.**
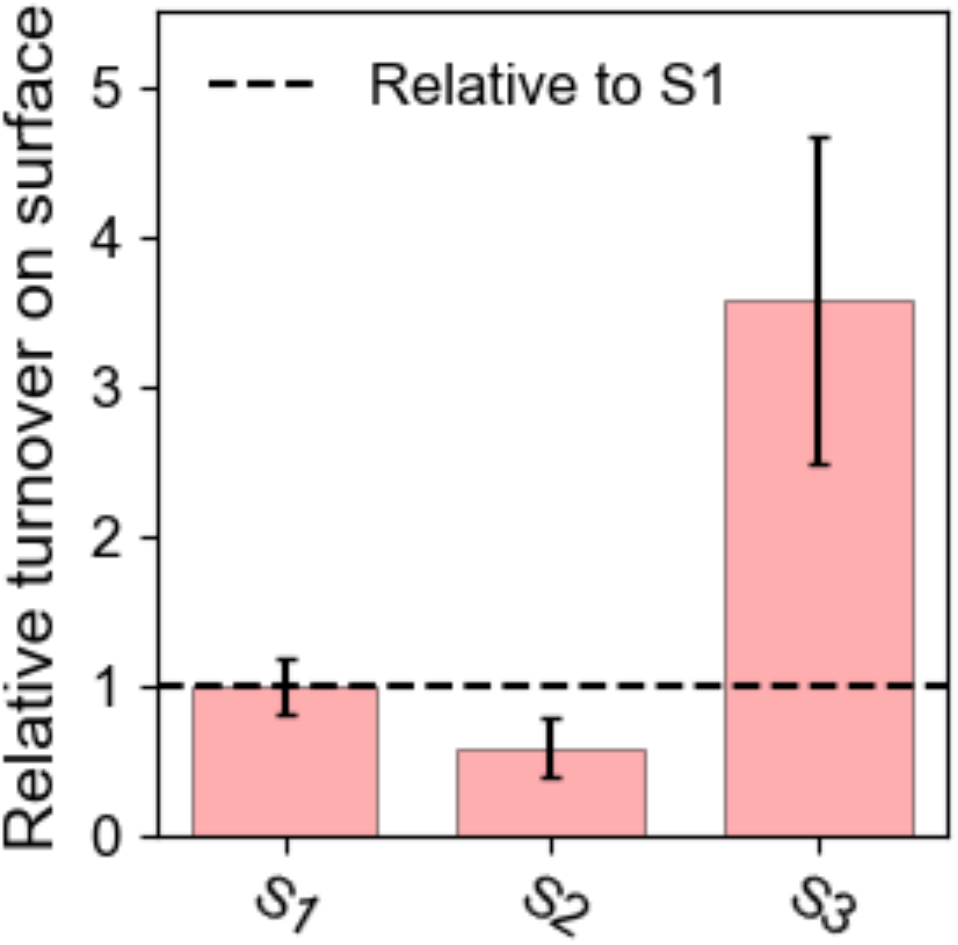
Relative turnover of Savinase variants. Comparison of relative activity per time, i.e., turnover, between variants shows S3 has 3.6 and 6.1 times higher turnover than S1 and S2, respectively. Turnover is calculated as the activity (Figure 2A) divided by the sum of the duration of binding events on the surface for S1, S2 and S3. Error bars correspond to the standard deviation of three technical replicates using error propagation.

**Figure 5.**
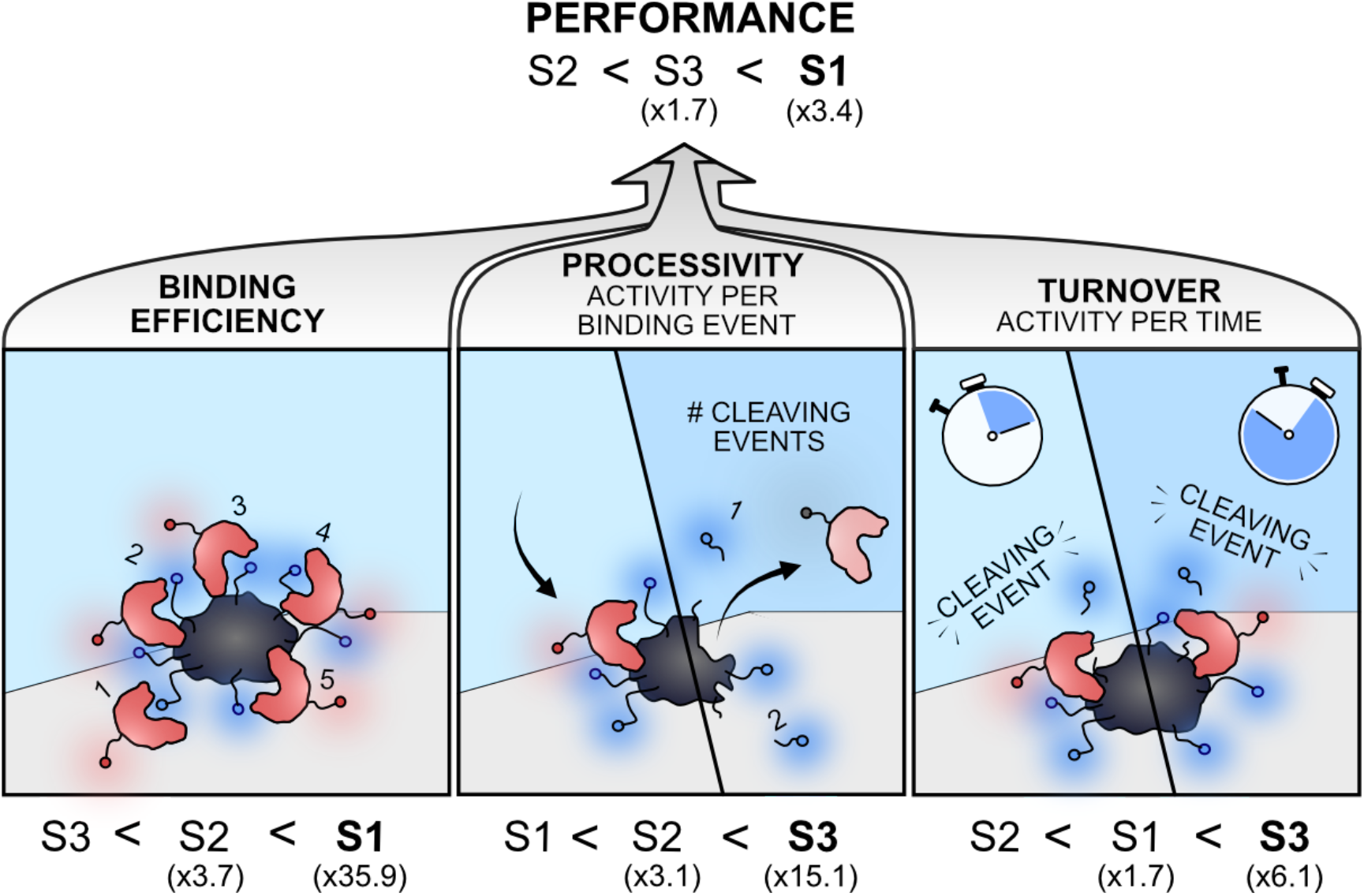
Components of proteolytic performance. Deconvolution of proteolytic performance of three Savinase variants into three elements 1) binding efficiency, 2) processivity and 3) turnover. Where S1 shows superior performance and binding efficiency, S3 demonstrates the highest processivity and turnover. S2 displays mediocrity on all three elements, resulting in the lowest overall performance.

Our results show that S1, the superior performer in the set, more than compensates for its relatively low processivity (Figure 2C) and intermediate turnover (Figure 4) by a superior binding efficiency (Figure 2B). S2 shows the lowest hydrolytic performance, with intermediate binding efficiency and processivity, along with the lowest turnover rate. Interestingly, the intermediate performance of S3 emanates from its superior processivity and turnover, albeit it having the lowest binding efficiency of the variants. In summary, in the case of S2 the mutations decreased both the binding efficiency and turnover on casein substrate as compared to S1. However, for S3, the increase in processivity and turnover partially compensates for its much-decreased binding efficiency, ultimately surpassing S2 but not S1 in performance.

## CONCLUSION AND OUTLOOK

Proteases are central proteins in numerous biological processes and applications. However, we have limited understanding of how their performance relates to molecular mechanisms as this layer of information can only be indirectly inferred from conventional ensemble assays. Here, we introduced a SM modality accomplishing direct observation of the binding dynamics of proteases to a proteinaceous substrate matrix with simultaneous monitoring of substrate degradation. The SM approach allowed us to deconvolute how the proteolytic performance of the protease Savinase interacting with casein micelles relies on binding efficiency, processivity and turnover number.

We investigated three variants of the protease, chosen for their varying performance in degrading casein. S1, the wild-type and highest performer in this system, excelled in terms of binding efficiency by far exhibiting the highest number of binding events. However, while variant S2 in turn bound more efficiently than S3, variant S3 showed the higher performance of these two. It follows that the activity per binding event, i.e. the number of cuts, must be different among the variants, indicating processivity. Analysis of the residence time distributions revealed three distinct characteristic binding states that were present across all three variants, but with varying sampling and lifetimes, consistent with previous studies proposing enzymes can sample multiple activity states^34,36,48–51^. Interestingly, evidence for conformation heterogeneity in Savinase in the substrate-interacting part of the protease has been reported^20,21^, which could be linked to the observed binding states. The binding events display second-scale dwell times, with up to ca. 20% of the events are binding with a characteristic time of 3 seconds or more. The extended nature of the interactions supports processivity. Additionally, it is evident that the three variants display significantly different turnover numbers when bound to the substrate. Interestingly, the inferior binding variant, S3, surpasses both S1 and S2 in this parameter to such an extent that it outperforms S2 in overall casein degradation performance. Conceivably, a path to a higher performing Savinase, in the context of the casein system, could be to pursue engineering of a high turnover number into a variant with more efficient substrate binding.

Understanding how differing substrate association and turnover rates influence protease performance offers insights for rational design and optimization. SM approaches emerge as powerful tools in molecular application understanding of enzymes and we envision that research utilizing the presented assay, or analogue SM techniques, can provide key insights required for the rational engineering of tailored enzymes involved in a host of industrial and biotechnological processes.

## METHODS AND MATERIALS

### Materials

SeTau-647-maleimide was purchased from SETA BioMedicals. Casein substrate (CAS: 9005-46-3) was bought from Merck and EnzChek casein BODIPY from Thermo Fisher Scientific. Fluorescent label Alexa Fluor 488 NHS Ester (Succinimidyl Ester) was purchased from Thermo Fisher Scientific. Chemical Trizma base (TRIS) (CAS: 77-86-1) was purchased by Sigma Aldrich, Denmark.

### Protease purification

Savinase variants were expressed in B. subtilis. Culture broth (CB) was added 3M Tris (1.6mL pr. 100mL CB) and GC850 (1.4mL pr. 100mL CB) before being spun down at 14.000 rpm for 20 minutes and filtered through a 0.2 µm filtration unit (Nalgene) to remove the host cells. The supernatant was applied to a MEP-hypercel (Pall) column and equilibrated in 20 mM Tris/HCl, 1mM CaCl_2_, pH 8.0. After washing the column with the equilibration buffer, the column was eluted with 20 mM AcOH/NaOH, 1mM CaCl_2_ pH 4.5. The eluted peak, containing the Savinase, was loaded onto an SP-Sepharose FF column (Cytiva), that had been equilibrated in 20 mm MES/NaOH, pH 6.0. The column was washed with the equilibration buffer and eluted with a linear gradient (0 to 0.5M) over three column volumes. The Savinase variants which eluted as a sharp peak were collected, and the purity was analyzed by Caliper LabChip (Revity); The major band observed belonged to the Savinase.

### Protease Fluorescent Labeling and Characterization

S1, S2 and S3 with single cysteine mutation at the C135 site, placed on the opposite side of the active site, were labeled with SeTau647-malemide, following previous protocols for enzyme labeling^35^. Labeling was characterized using intact mass electrospray ionization mass spectrometry (ESI-MS) (Figure S10) and labeling efficiencies were determined using NanoDrop (Table S3). Comparing the wild-type with the single cysteine mutation S1, showed little to no difference in activity (Figure S11).

### Substrate Fluorescent Labeling and Characterization

The substrate, casein, was labeled with Alexa Fluor 488 NHS Ester. 1 μL of 10 mg/mL Alexa Fluor 488 NHS Ester diluted in DMSO was added to 1500 μL 10 mg/mL casein in PBS pH 8.3 buffer. The sample was left to incubate at room temperature for two hours. Hereafter, the sample was spin-filtered through a 3 kDa filter until the filtered liquid had no color. The remaining labeled protein was spun out of the filter and measured using NanoDrop to determine the concentration and the labeling efficiency of 91%.

### Substrate Surface Preparation

A 6 well ibidi sticky-Slide VI 0.4 was attached to a dried and clean microscope glass slide. 80 *µ*L of 6 nM casein substrate in 50 mM TRIS pH 8 solution was added to each of the 6 wells and incubated for a minimum of 15 minutes, where adsorption ensured immobilization of casein micelles on the surface. Hereafter, each of the wells were washed 5 times with buffer before imaging.

### TIRF Microscopy

TIRF microscope (IX 83, Olympus) was used for the SM experiments. We utilized an oil immersion objective (UAPON 100XOTIRF, Olympus) in conjunction with an EMCCD camera (ImageEM X2, Hamamatsu), resulting in a pixel width of 160 nm. The laser lines of 488 nm and 640 nm were used to excite the fluorophores Alexa Fluor 488 (casein substrate) and SeTau647 (Savinase variants), respectively. Imaging was done with an exposure time of 100 ms, a penetration depth of 100 nm and 300 EM gain in the streaming setting, resulting in a frame length of 101 ms. Videos were recorded with a laser power of 8.4% and 10% for the 488 nm laser line and 640 nm laser line respectively. After the addition of 0.05 nM Savinase (S1, S2 or S3), in TRIS pH 8 solution, in the given well, one frame was recorded in the substrate channel (488 nm) for every 1000 frames in the enzyme channel (640 nm), repeated four times, totalling approximately 7 minutes per experiment. Only one experiment was conducted per well in the prepared 6 well experiment slide, which was started immediately after the addition of proteases.

### Ensemble Activity Assay

Ensemble activity measurements were done on a SpectraMax iD3 Multi-Mode Microplate Reader. Here, a concentration of 0.7 *µ*M casein BODIPY was used with a concentration of 0.1 *µ*M Savinase. The measurement was started immediately after the addition of Savinase to the chambers containing casein. Fluorescence was recorded using an excitation wavelength of 590 nm and an emission wavelength of 625 nm. All measurements were repeated thrice.

### Extraction of SM and Ensemble Performance

SM performance was calculated as the reciprocal to the relative decrease in intensity of the TIRF field of view in the casein channel for each experiment. The ensemble performance was found as the slope coefficient of the first 2.5 minutes of measurements on the Microplate Reader (for elaboration see Figure S4).

### Image Analysis and SPT

For localization and tracking of individual Savinase molecules over time, an in-house algorithm based on TrackPy^52^ was used. The script localized each molecule using an intensity threshold based on the signal-to-noise-ratio (SNR) of the given video. Hereafter, the SM detections were linked through time using the nearest neighbor algorithm incorporated in TrackPy^52^.

### Extraction of Residence Times

The residence time of each trace was extracted from the number of frames a molecule present in the SPT detections (see Figure S6). To minimize shot noise from the dataset, events that lasted for a single 100 ms frame, were excluded.

### Normalization

All displayed bar plots are normalized to S1 for easy comparison, this is done by dividing a given raw value for each of the variants, e.g. performance, number of binding events, occupancy of binding states etc., with the raw value for the given feature for S1. Normalized errors were determined using error propagation.

### Resolving Binding States

The distributions of the extracted residence times of Savinase variants were fitted using unbinned MLE, determining a three binding state model using a triple exponential distribution (equation 1) best described the binding behavior of Savinase variants. The fitting parameters were found iteratively until convergence of the likelihood function to find the optimal parameters and fit.

## ASSOCIATED CONTENT

### Supporting Information

Tracking statistics, labeling characteristics and efficiencies of Savinase variants and substrate, methodology for extraction of relative activity of Savinase variants for SM and ensemble measurements, activity and residence times of Savinase for entire field of view versus only for detected casein micelles, control experiments of activity and binding behavior of Savinase without casein substrate, probability of sampling each of the three extracted binding states for the three Savinase variants given a specific residence time, evidence of multiple fluorescent labels per casein micelle, evidence of the existence of a casein carpet covering the entire surface at the experimental condition (<u>PDF</u>)

## AUTHOR INFORMATION

### Author Contributions

Conceptualization, S.M.C. and N.S.H.; methodology, E.W.S. and N.S.H.; software, E.W.S, A.F., S.B. and J.K.H.; validation, E.W.S., F.B.R. and N.S.H.; formal analysis, P.H. and E.W.S.; investigation, E.W.S., F.B.R, J.S.L. and E.V.; resources, S.M.C. and N.S.H.; data curation, E.W.S., F.B.R. and J.S.L.; writing—original draft preparation, E.W.S.; writing—review and editing, E.W.S., J.K.H., S.M.C. and N.S.H.; project administration, N.S.H.; overall project management and supervision, N.S.H.; funding acquisition, N.S.H.. All authors have read and agreed to the published version of the manuscript.

## Notes

Sune Christensen and Jeannette de Sparra Lundin are employed by Novonesis.

## ACKNOWLEDGMENT

The authors give thanks to Gustav Hammerich Hansen for assistance with protein purification. This work was supported by the Villum foundation center BIONEC (18333), the NNF challenge center for Optimised Oligo escape, the NNF center for 4D cellular dynamics, the Lundbeck foundation grant R346-2020-1759, Villum foundation Synergy grant (40578), and Carlsberg foundation grant CF21-0499. NSH is a member of the Integrative Structural Biology Cluster (ISBUC) at the University of Copenhagen and associate member of the Novo Nordisk Foundation Center for Protein Research, which is supported financially by the Novo Nordisk Foundation (NNF14CC0001).

## ABBREVIATIONS

SPT: single particle tracking
SM: single-molecule
MLE: maximum likelihood estimation
SNR: signal-to-noise-ratio
TIRF: total internal reflection fluorescence
S1: Savinase 1
S2: Savinase 2
S3: Savinase 3
ROI: region of interest
ESI-MS: intact mass electrospray ionization mass spectrometry

