## Supplementary information for "Proteolytic Performance is Dependent on Binding Efficiency, Processivity and Turnover: Single Protease Insights"

^†^ Novonesis, Biologiens Vej 2, DK-2800, Kgs. Lyngby, Denmark.

**Corresponding Authors**

* (N.S.H.).

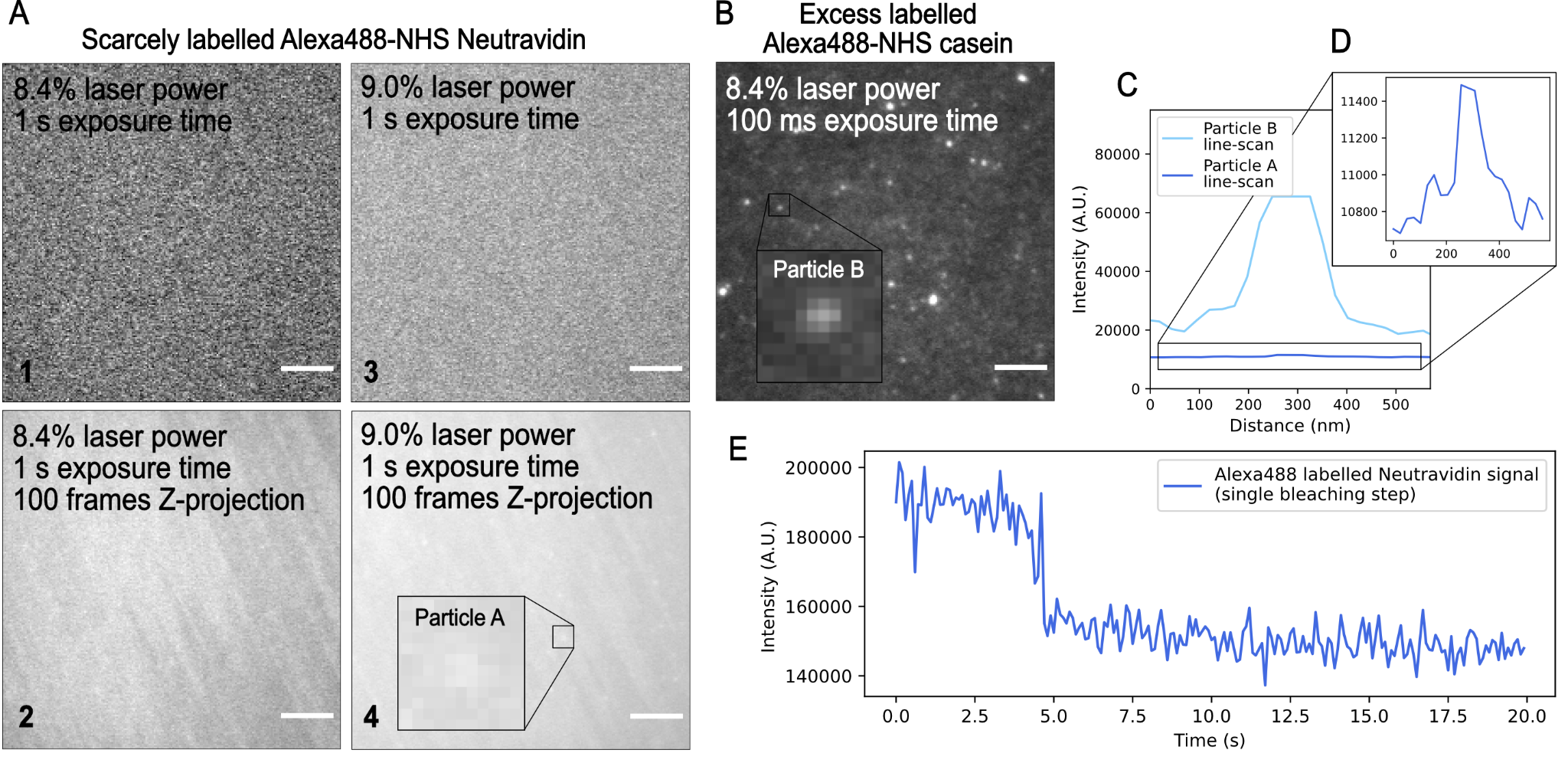
**Figure S1.** **Scarcely labeled Neutravidin with Alexa488 NHS ester versus excess labeled casein with Alexa488, showing casein micelles having multiple fluorophores.** A) TIRF images of PLL-g-PEG-biotin covered surface after addition of Alexa488 NHS ester labeled Neutravidin followed by washing, for 1) 8.4% laser power, the laser power used for the main experiments with casein, and 1 second exposure time, 2) the Z-projection of the average intensity of 100 frames of 1), showing no clear particle signals, 3) 9.0% laser power and 1 second exposure time and 4) the Z-projection of the average intensity of 100 frames of 3), showing vague particle signals, as such a single fluorophore is barely visible at this intensity. Zoom in of a particle, single Alexa488 labeled Neutravidin, Particle A. B) TIRF image of 6 nM of Alexa488 NHS ester labeled casein at 8.4% laser power and 100 ms exposure time, showing clearly bright particles. Zoom in of a particle, comparatively small casein micelle, Particle B. C) Line-scan of Particle A (single fluorophore), zoom in D), versus line-scan of Particle B (small casein micelle), clearly showing that even at a lower laser power (8.4%), the casein signal, even the background, is higher than the single fluorophore at laser power 9.0%. Line-scans were extracted using ImageJ^1^. E) Raw intensity over time of Particle A, at a laser power of 80% and 100 ms exposure time, showing a single bleaching step, confirming that this is a single fluorophore. Extracted using in-house tracking algorithm based on TrackPy^2^, see Methods and Materials.

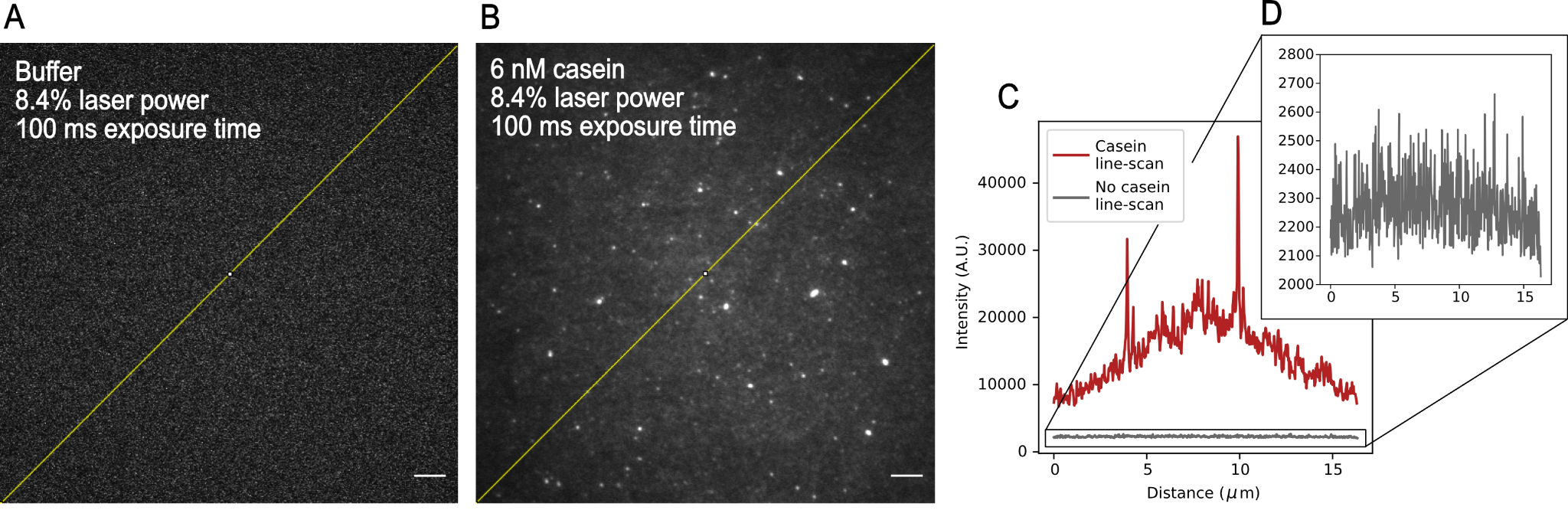
**Figure S2.** **Surface intensity with and without Alexa488 NHS ester labeled casein at same instrument settings.** A) Raw image of clean surface in buffer at 8.4% laser power and 100 ms exposure time, displaying line-scan as shown in C/D), B) Raw image of 6 nM casein surface in buffer at 8.4% laser power and 100 ms exposure time, displaying line-scan as shown in C). C) Raw intensity line-scan extracted from ImageJ^1^ of A), no casein gray trace, and B), 6 nM casein red trace, and D) zoom in of no casein line-scan, displaying clear intensity difference and the existence of a casein carpet as even the “background” intensity and the lowest displayed intensity is significantly higher than when no casein is on the surface. This confirms that the entire surface is covered in fluorescently labeled casein at a concentration of 6 nM. Note that neither images were background corrected prior to taking the line scan, as such the intensity peaks in the middle of the line-scan, to ensure comparable intensities.

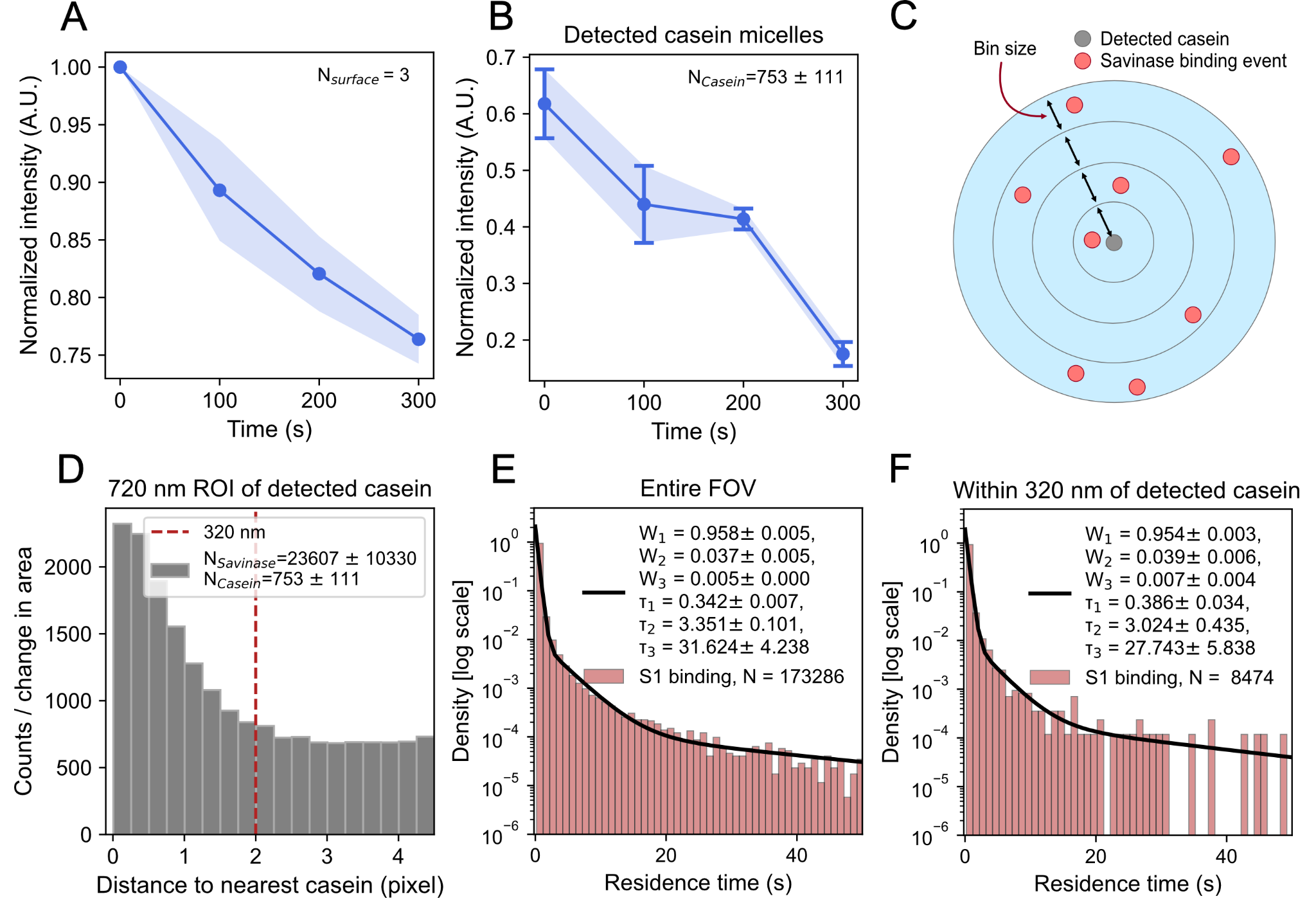
**Figure S3.** **Representative casein surface and binding behavior of Savinase across the entire field of view (FOV) and only within a region of interest (ROI) of detected casein particles.** A) Normalized summed intensity over time of entire FOV in casein channel after addition of S1, showing one and two standard deviations. B) Normalized average intensity of all detected casein micelles. Here, the detected casein micelles have a higher concentration of protein than the surrounding carpet, detections done as outlined in Methods and Materials, in FOV in casein channel over time after addition of S1, for the exact same experiment as A), showing one and two standard deviations. Here, a larger relative intensity decrease is expectedly observed, as compared to A), as we are looking at more concentrated areas of substrate which will experience more degradation than lower concentration areas. C) Representation of observed distance between locations of large/detected casein micelles and S1 binding events, showing a possible binning of distances as shown in D). D) Number of all detections within 800 nm of a detected casein micelle, measured per square nanometer to normalize to increasing area as distance to the center position of a casein micelle increases and shown in C). Red line indicates the cut-off point at 400 nm from nearest casein micelle, where the distribution starts to even out to a background level. E) Residence time distribution of S1 binding events on the entire FOV on the casein surface, with triple exponential fit and the standard deviation of each of the fit parameters. F) All durations of S1 binding events within 400 nm of a detected casein micelle on casein surface, with triple exponential fit and the standard deviation of each of the fit parameters, which show to be very similar to the fit for all binding events. All plots and uncertainties based on four replicates of experiments conducted at pH 8 with S1.

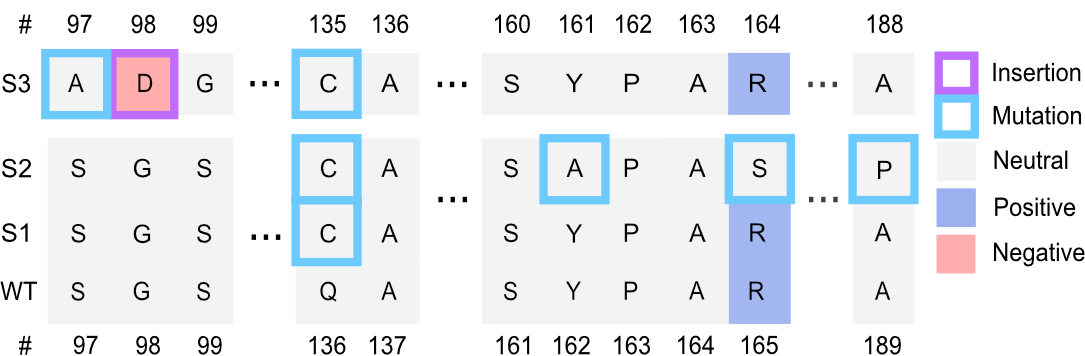

**Figure S4. Sequence alignment of the S1, S2 and S3**, color coding denotes the approximated charge of the amino acids in Savinase structure at pH 8, calculated using propka3.5.0^3^. The mutation to S2 and S3 all reside in the periphery of the cleft down to the active site. S2 and S3 have a more negative net charge than S1.

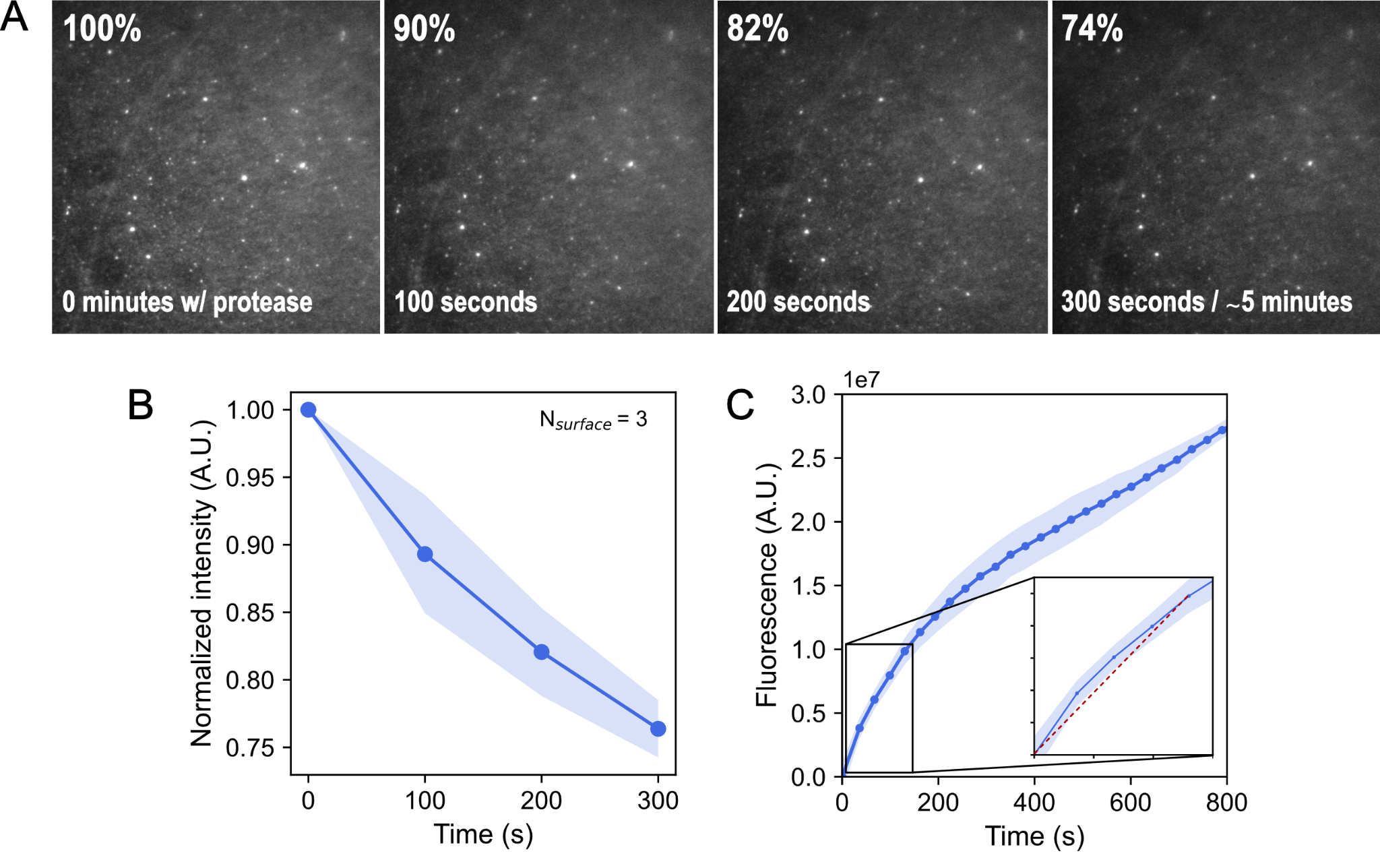
**Figure S5.** **Analysis of activity measurements of Savinase by ensemble and SM assay.** A) Raw images of decreasing intensity of field of view of TIRF images in casein channel after addition of Savinase at pH 8, shows the percentage decrease over time of the summed intensity of entire field of view. B) Representative plot of normalized sum of intensity in casein channel over time, showing initial and final intensity of experiment, where the reciprocal of the final intensity was used as the measure of SM activity. C) Fluorescence plate reader measurement of casein BODIPY after addition of Savinase and zoom in of linear slope/rate, the first 2.5 minutes, equating to the utilized ensemble activity measure of Savinase.

| **Substrate,**  **Condition** | **Savinase variant** | **Measurement** | **Repetitions** | **Detections /**  **per experiment** |
| --- | --- | --- | --- | --- |
| Casein /  Buffer | Savinase 1 | Plate-reader (Ensemble) | 3 / 3 | --- |
|  | Savinase 2 |  |  |  |
|  | Savinase 3 |  |  |  |
|  | Savinase 1 | TIRF (Single molecule) | 3 / 4 | 57762 ± 19527 /  1303 ± 247 |
|  | Savinase 2 |  |  | 5368 ± 2878 /  740 ± 357 |
|  | Savinase 3 |  |  | 2240 ± 277 /  392 ± 148 |

**Table S1. Number of repetitions per experimental condition**, and average number of SM detections, by TIRF microscopy, for each Savinase variant, S1, S2 and S3. Errors correspond to the standard deviation of the three or four technical replicates as indicated in the repetition column.

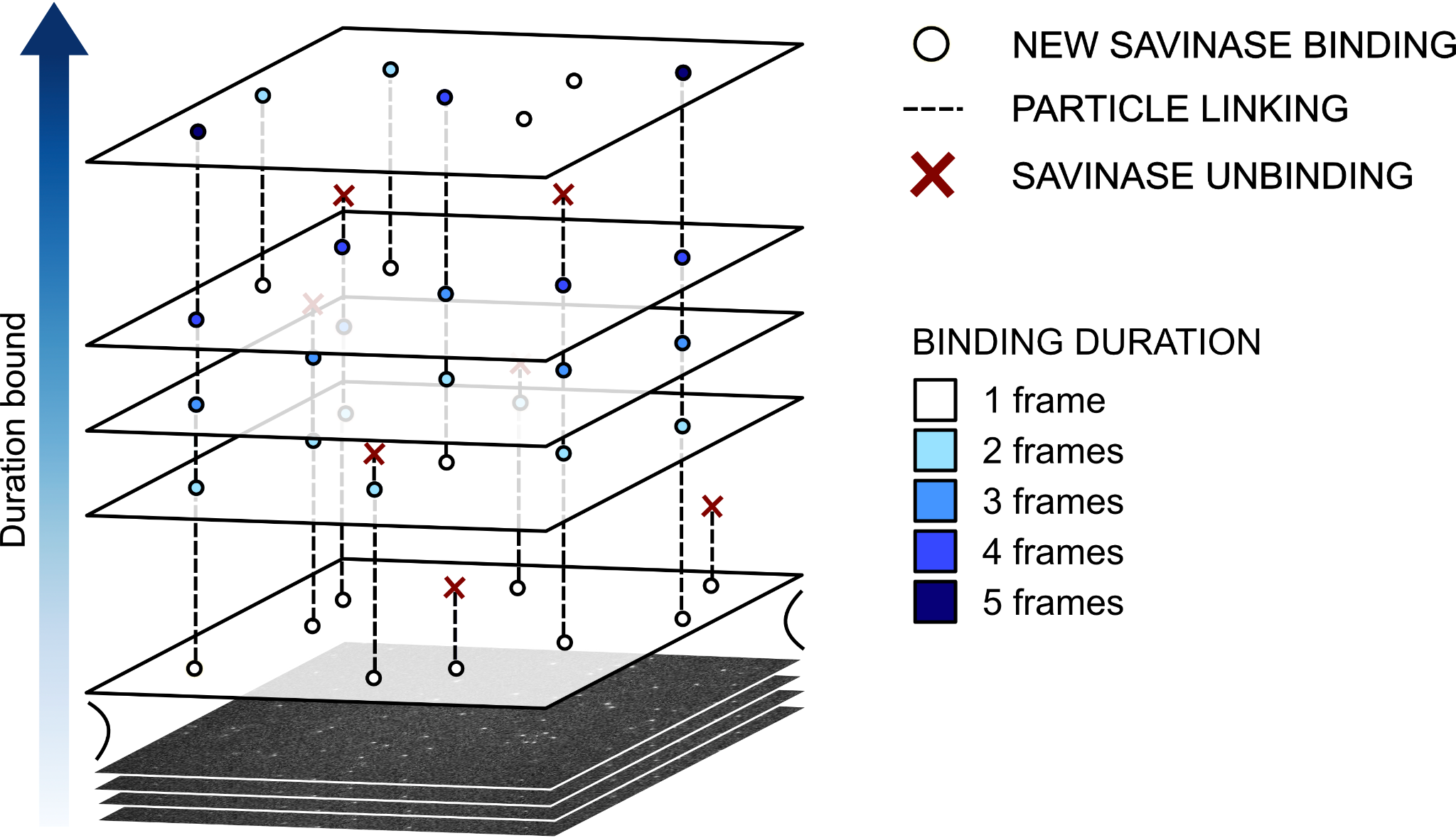

**Figure S6.** **Illustration of parallelized recordings of multiple reversible binding events of Savinase on casein covered surface**. Dotted lines indicate binding for multiple frames and red X indicates discontinued tracks due to unbinding. Binding duration of transient events were determined based on the number of frames the molecule was observed by the utilized tracking software (see Methods and Materials).

**Equation S1. Calculation of probability of binding state at specific residence time.**

For residence time t, the probability of the protease being in a given state x (1: short-lived, 2: intermediate-lived or 3: long-lived) is given by:

$$P_{x}(t)=\frac{\frac{W_{x}}{\tau_{x}}\times exp\left( -\frac{t}{\tau_{x}} \right)}{\frac{W_{1}}{\tau_{1}}\times exp\left( -\frac{t}{\tau_{1}} \right) + \frac{W_{2}}{\tau_{2}}\times exp\left( -\frac{t}{\tau_{2}} \right) +\frac{W_{3}}{\tau_{3}}\times exp\left( -\frac{t}{\tau_{3}} \right)}$$

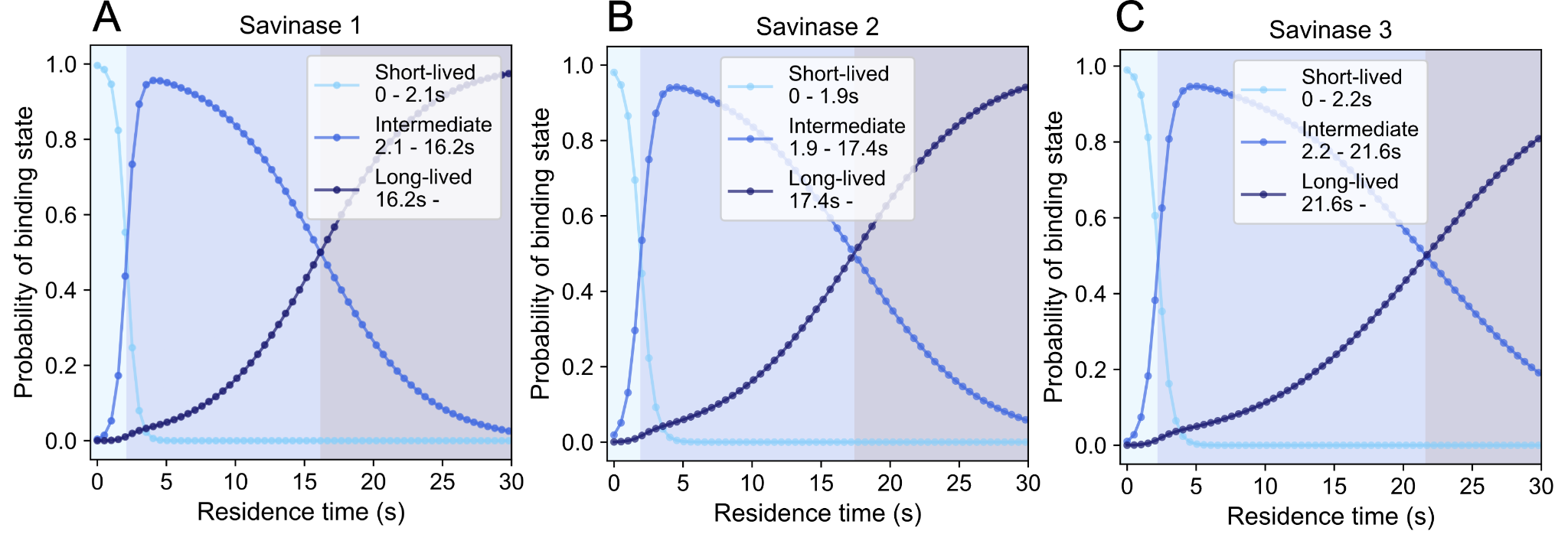
**Figure S7. Probability of binding state given specific residence time.** Probability, from unbinned maximum likelihood fitting with triple exponential function (equation 1 in main article) using Equation S1, of a given residence time representing a given binding state based on maximum likelihood estimation fitting for A) S1 showing the kiss and run state to be most likely at residence times from 0-2.1 seconds, the short-lived from 2.1-16.2 seconds and the long-lived binding at residence times longer than 16.2 seconds, B) S2 showing the kiss and run state to be most likely at residence times from 0-1.9 seconds, the short-lived from 1.9-17.4 seconds and the long-lived binding at residence times longer than 17.4 seconds and C) S3 showing the kiss and run state to be most likely at residence times from 0-2.2 seconds, the short-lived from 2.2-21.6 seconds and the long-lived binding at residence times longer than 21.6 seconds. The ranges for the most likely states are determined based on the interception of probability functions.

|  | **S1 w/ casein** | **S2 w/ casein** | **S3 w/ casein** |
| --- | --- | --- | --- |
| **Total time spent in binding states (seconds) / experiment** | | | |
| **𝝉1** | 19246 ± 6479 | 1953 ± 938 | 897 ± 94 |
| **𝝉2** | 7157 ± 2044 | 3371 ± 1629 | 1218 ± 40 |
| **𝝉3** | 8210 ± 1954 | 11784 ± 10990 | 3535 ± 1461 |

**Table S2. Sum of time spent in each binding for Savinase variants** S1, S2 and S3 on casein substrate per experiment. Determined by summing all residence times of ranges calculated and shown in Equation S1 and Figure S7.

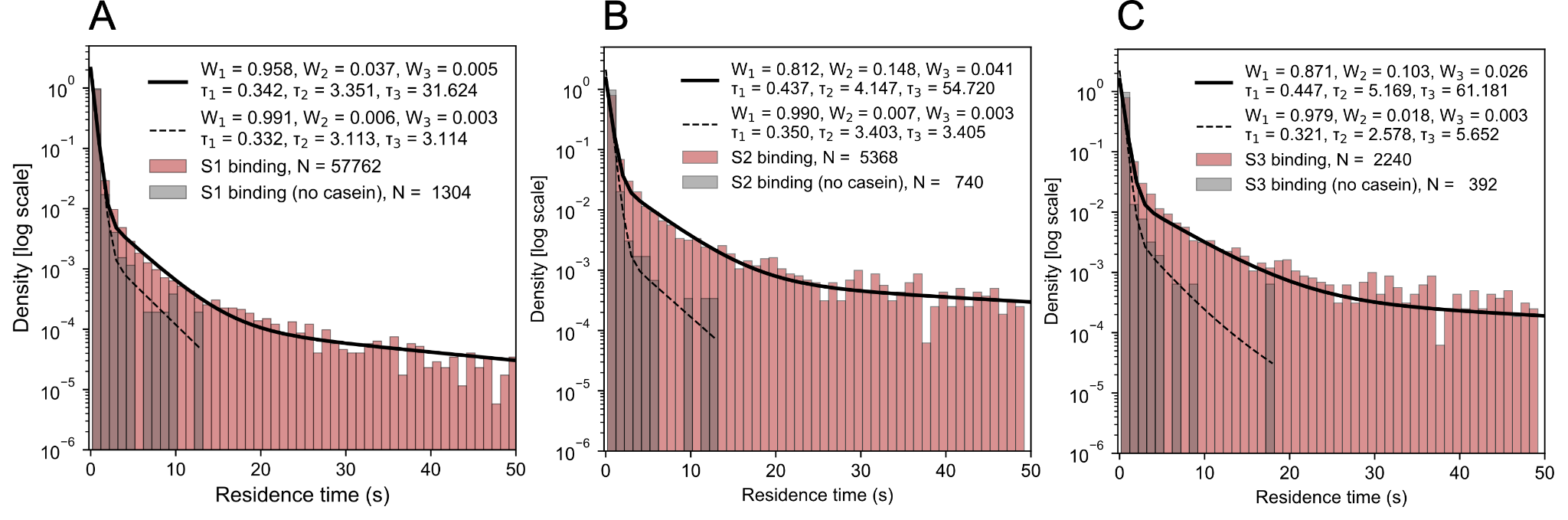
**Figure S8. Distribution of residence times with and without substrate present for** A) S1, B) S2 and C) S3. MLE fitting reveals that the parameters *τ*_2_ and *τ*_3_ converge to the same value, namely, the intermediate-lived state, *τ*_2_, when substrate is present (Figure 3C and S7). This suggests that sampling of the third and long-lived binding state occurs explicitly in the presence of substrate. For S3 the long-lived state, *τ*_3_, is found to be slightly different from the short-lived state, *τ*_2_, when no casein is present, however, these are still quite similar and averages around the value *τ*_2_ when casein is present.

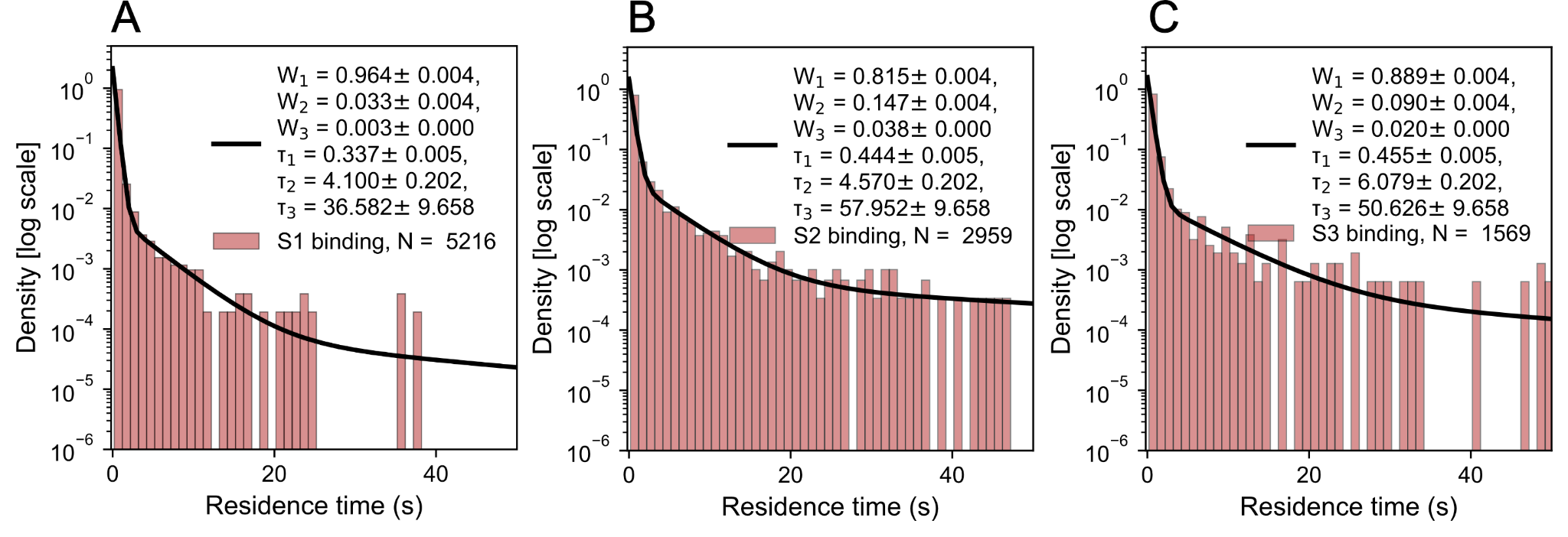
**Figure S9.** **Random sampling of residence times on substrate for Savinase variants.** Each dataset of residence times for the Savinase variants on casein is sampled randomly (using the random package for Python) the same number of times as there were binding events on a non-casein surface to demonstrate that sampling of the long-lived binding state happens even with this fewer binding events for A) S1, B) S2 and C) S3.

|  | **Savinase 1** | **Savinase 2** | **Savinase 3** | **Casein** |
| --- | --- | --- | --- | --- |
| Labeling efficiency (%) | 53.2 | 53.3 | 53.4 | 91.1 |

**Table S3. Labeling efficiency of utilized Savinase variants**, Savinase 1 (S1), Savinase 2 (S2) and Savinase 3 (S3), and casein substrate, determined by NanoDrop measurements.

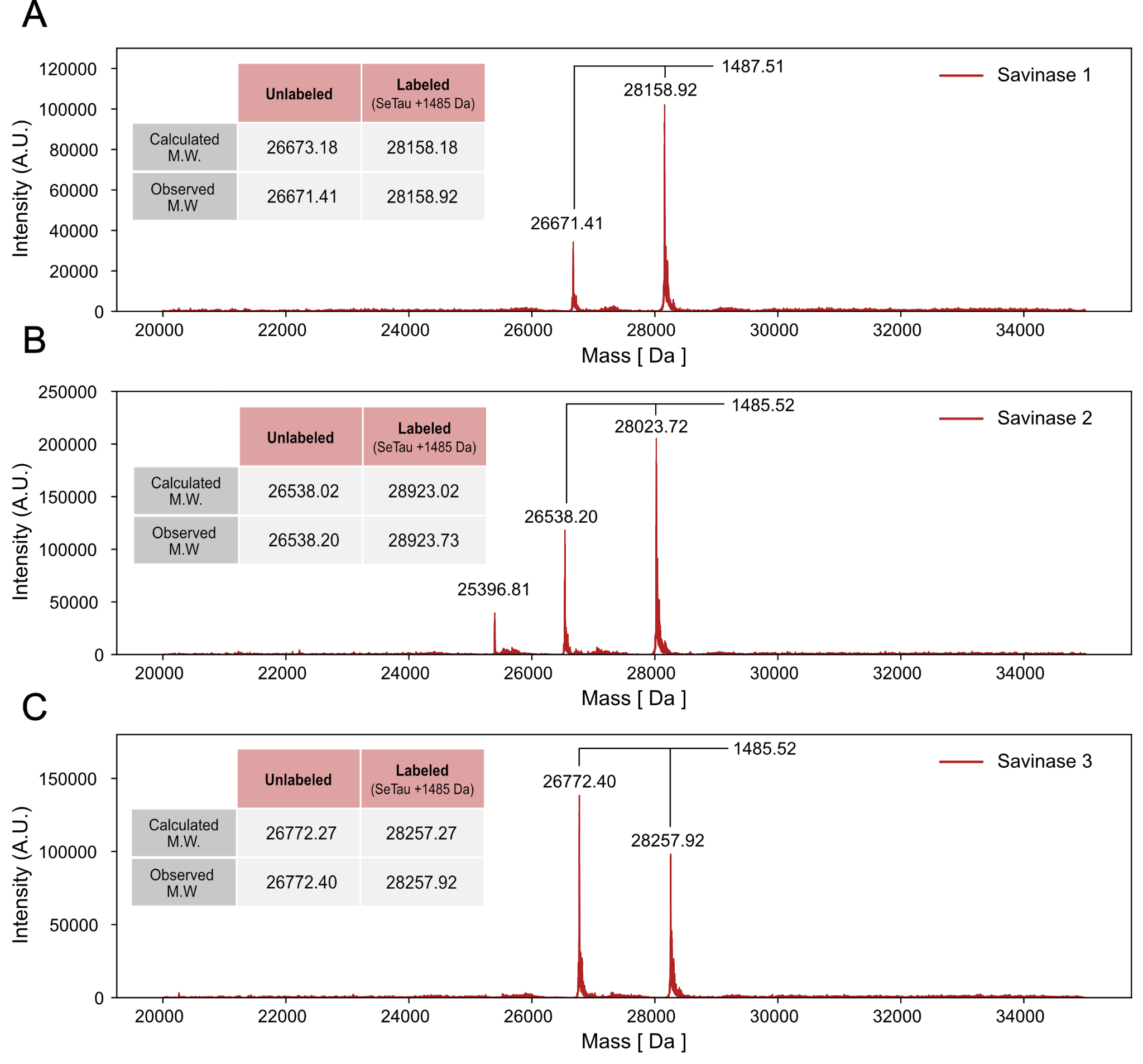
**Figure S10. Mass spectrometry characterization** confirmed fluorescent labeling of Savinase variants with SeTau647 for A) S1, B) S2 and C) S3.

**Figure S11.** **Effect of single cysteine mutation on proteolytic activity.** Comparison of ensemble proteolytic activity, measured by Microplate Reader on casein substrate as explained in Methods and Materials, for unlabeled S1 with, gray trace, and without, red trace, single cysteine mutation necessary for singular fluorescent labeling. Little to no effect of mutation is observed.
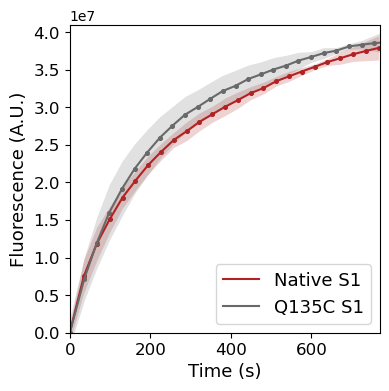

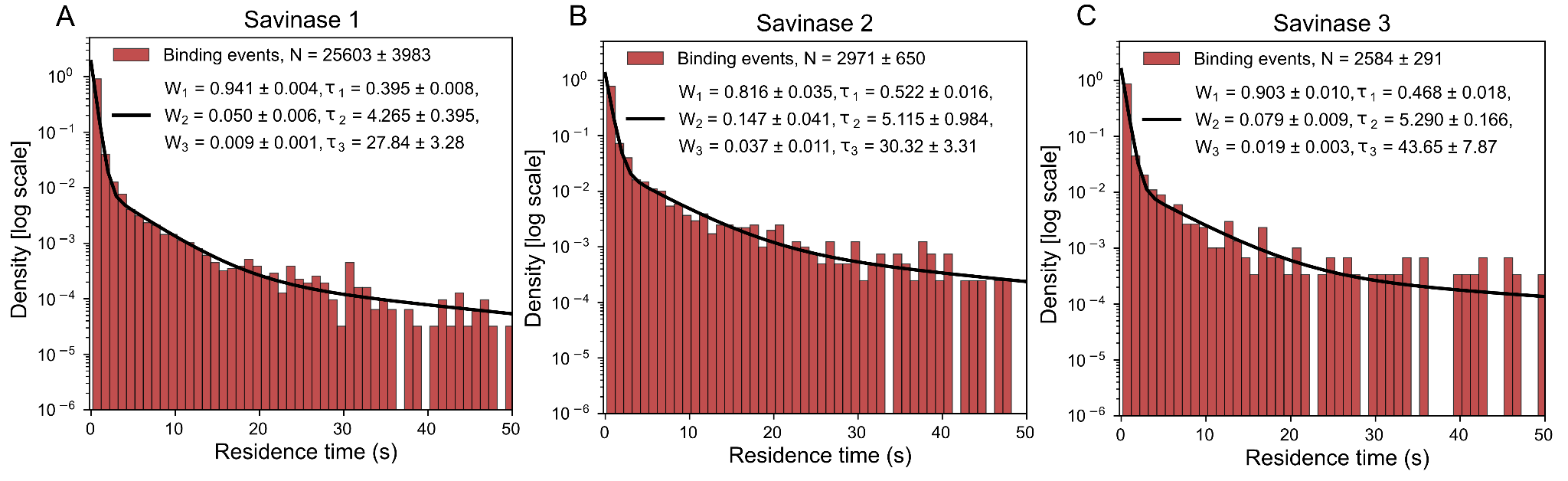

**Figure S12. Savinase variants binding behavior and different preparation of casein substrate** (as described in Methods and Materials), shows moderately different occupancies and lifetimes of binding states. This is to be expected for a proteinaceous substrate matrix of different constituents^4^, due to slight variation in composition affecting the electrostatic interactions between protease and substrate.
